# Engineering Phage Host-Range and Suppressing Bacterial Resistance Through Phage Tail Fiber Mutagenesis

**DOI:** 10.1101/699090

**Authors:** Kevin Yehl, Sébastien Lemire, Andrew C. Yang, Hiroki Ando, Mark Mimee, Marcelo Der Torossian Torres, Cesar de la Fuente-Nunez, Timothy K. Lu

## Abstract

The rapid emergence of antibiotic-resistant infections is prompting increased interest in phage-based antimicrobials. However, acquisition of resistance by bacteria is a major issue in the successful development of phage therapies. Through natural evolution and structural modeling, we identified host-range determining regions (HRDR) in the T3 phage tail fiber protein and developed a high-throughput strategy to genetically engineer these regions through site-directed mutagenesis. Inspired by antibody specificity engineering, this approach generates deep functional diversity (>10^7^ different members), while minimizing disruptions to the overall protein structure, resulting in synthetic “phagebodies”. We showed that mutating HRDRs yields phagebodies with altered host-ranges. Select phagebodies enable long-term suppression of bacterial growth by preventing the appearance of resistance in vitro and are functional in vivo using a mouse skin infection model. We anticipate this approach may facilitate the creation of next-generation antimicrobials that slow resistance development and could be extended to other viral scaffolds for a broad range of applications.

**Highlights:**

- Vastly diverse phagebody libraries containing 10^7^ different members were created.
- Structure-informed engineering of viral tail fibers efficiently generated host-range alterations.
- Phagebodies prevented the development of bacterial resistance across long timescales *in vitro* and are functional *in vivo*.

## INTRODUCTION

The rapid escalation of drug-resistant bacterial infections and under-resourced antibiotic pipeline make it imperative to develop alternative therapies. A resurging approach gaining significant interest is phage therapy, in which bacteriophages are used as antimicrobials (Bikard et al., 2014; Chen et al., 2014; Citorik et al., 2014; Devlin et al., 2016; Kutateladze and Adamia, 2010; Kutter et al., 2010, 2015; Lu and Collins, 2007, 2009; Maynard et al., 2010; Shen et al., 2015). Because bacteriophages are functionally orthogonal to antibiotics, they are generally unaffected by antibiotic-resistance mechanisms, making them promising for treating such infections (Międzybrodzki et al., 2012). Moreover, phage therapy has successfully been used for compassionate care cases when all antibiotic treatment options have failed (Dedrick et al., 2019; Schooley et al., 2017). In addition, phages are selective for particular bacterial strains, as opposed conventional chemical antibiotics that exhibit broad-spectrum activity, which contributes to antibiotic-resistance development. Phage selectivity is dependent on binding to cell surface receptors in order to recognize their host and initiate infection (Silva et al., 2016). However, reliance on receptor recognition for infectivity implies that resistance against a bacteriophage can occur through receptor mutations. As a result, phage-based products are usually composed of multiple unrelated phages that collectively target a range of receptors, thus distributing the selective pressure away from any individual phage receptor. However, these cocktails are often composed of uncharacterized phages whose biology are poorly defined. Furthermore, manufacturing cocktails composed of diverse phages, as well as tracking their pharmacodynamics and immunogenic properties, is complex, which can limit their development as antimicrobial drugs (Cooper et al., 2016).

Various approaches have been undertaken to rationally expand the host-range of phages to combat resistance (Ando et al., 2015; Chen et al., 2017; Gebhart et al., 2017; Hawkins et al., 2008; Heilpern and Waldor, 2003; Lin et al., 2012; Nguyen et al., 2012; Scholl et al., 2009; Yoichi et al., 2005; Yosef et al., 2017). However, these approaches depend on hybridization between already characterized bacteriophages with known and desired host-ranges. Natural evolution of phages has also been harnessed to alter phage host-range, but the lack of control over where mutations occur can hamper our understanding of structure-to-function mapping. Therefore, the development of targeted and high-throughput methodologies to rapidly broaden phage host-range and to overcome bacterial resistance could pave the way for next-generation phage technologies.

To circumvent these technological barriers, we performed targeted mutagenesis on well-defined regions of the phage that are essential for host recognition, thus increasing the sequence space screened at these crucial loci. This approach is analogous to antibody specificity engineering, in which vast amounts of diversity are generated at particular epitope binding regions. Similar to phage tail fibers, antibodies are highly selective to their target antigen (Beck et al., 2010; Foltz et al., 2013). The epitope recognition region of antibodies lies at the tip of the interface formed by the light and heavy chains (Figure 1A) and is particularly directed by three hypervariable regions called complementary determining regions (CDR). Mutations in these regions alter the target specificity of the antibody (Ducancel and Muller, 2012; Igawa et al., 2011). Phage T7 tail fibers also display CDR-like loops at the tip of the tail fiber (Figure 1A), which inspired us to ask whether a phage’s host specificity can be manipulated through changes in these small regions. Motivated by the similarity between antibodies and phage tail fibers (Figure 1A), we developed a strategy to engineer the host-range of the phage T3, a T7-like phage.

**Figure 1:**
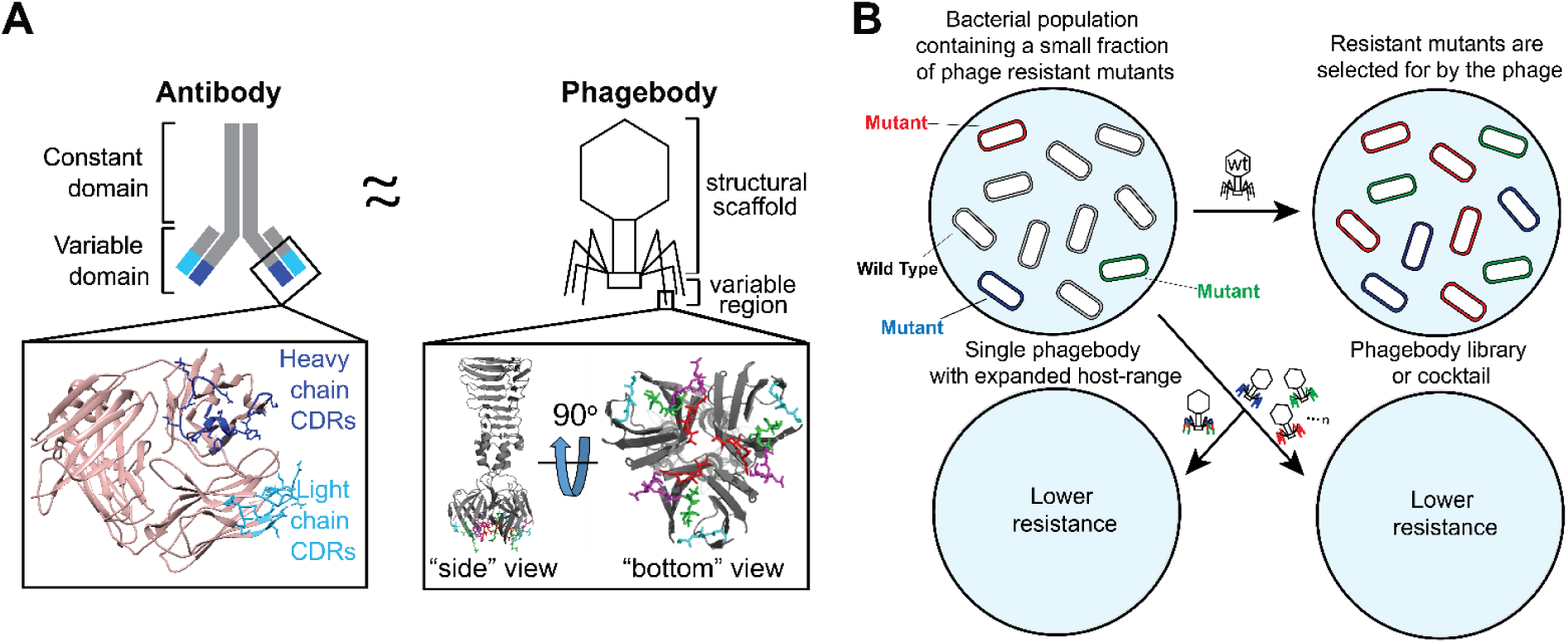
Phagebody design and proposed ability to target bacterial mutants. (**A**) Schematic illustrating the similarities between antibodies and the phagebody tail fiber. In an antibody, antigen recognition is primarily encoded by six hypervariable complement-determining regions (CDRs), three located on the heavy chain and three on the light chain. The inset (left) presents the three-dimensional structure of the variable domain of an antibody (PDB ID 1IGT). Heavy chain CDRs are colored dark blue and light chain CDRs are colored teal. In phage T7, host-range is largely determined by the C-terminus of its tail fiber protein, gp17. The inset (right) shows the crystallographic structure of the C-terminal 182 amino acids of T7 gp17 (PDB ID 4A0T). Outward loops (red, magenta, green, and light blue) are expected to participate in receptor recognition while tolerating mutations. Phagebodies are designed to carry mutations in these loops while leaving other structures of the tail fiber intact. (**B**) Schematic illustrating how resistance appears in bacterial cultures and how phagebody cocktails or individual phagebodies are proposed to suppress resistance.

Regions in the T3 tail fiber were targeted for mutagenesis to create highly diverse phage libraries that were screened to identify phages with altered host-ranges. We sought to expand the host-range of T3 to target naturally occurring phage-resistant bacterial mutants and delay or even prevent the onset of phage resistance (Figure 1B). We refer to our engineered phages as “phagebodies” because of the inspiration from antibody specificity engineering. Importantly, we discovered “hot spot” regions in the tail fiber of T3 that dictate phage host-range and generated phages that suppress bacterial resistance. We envision this strategy will enable high-throughput studies to elucidate phage-host interactions and support the production of phagebodies that effectively treat AMR infections. Furthermore, this work demonstrates the feasibility of developing approaches that directly address the evolution of bacterial resistance to antimicrobials, which is currently a major limitation for the successful development of new antimicrobial therapeutics.

## RESULTS

### Identification of host-range determining regions (HRDR) for T3

Evolution of phage T3 co-cultured with its host *E. coli* B proceeds through a limited number of pathways and is predictable (Perry et al., 2015). Bacteria first evolve resistance through acquiring mutations in genes that alter the LPS structure, which in turn select for T3 mutants that can infect these bacterial mutants. Ultimately, the phage mutants drive the selection for secondary mutants of *E. coli B*, which the phage are unable to prevent from overtaking the culture. Therefore, we used T3 resistance as a model system for validating our proposed engineering approach.

T3 initially binds to the bacterial LPS with its tail fibers for host recognition. Each tail fiber is composed of a homotrimer of the gene *17* product, gp17, which we have previously identified the carboxyl-terminal, ∼450-553 amino acid globular domain or the “tip”, as a determinant for host specificity (Ando et al., 2015). The structure of the tip region of T3 gp17 (residues 454-558) was generated via homology modeling using the published structure of T7 gp17 (PDB 4A0T) (Garcia-Doval and van Raaij, 2012) as a template (Swiss-model) (Arnold et al., 2006) (Figure 2A). This structure was used to identify potential host-determining regions for T3 (Figure 2B). The distal 104 a.a. portion of gp17 forms an intertwined globular domain shaped by an eight-stranded beta barrel (strands labeled B to I). The strands are connected by random coils, which are referred to by the beta strands they connect. Three of these coils, CD, EF and GH, are oriented towards the tail fiber shaft (“inward loops”) and are hypothesized to not partake in host recognition. The four other coils, BC, DE, FG and HI, are displayed on the opposite side of the beta barrel and point away from the tail fiber (Figure 2A and B; “outward loops”, highlighted in magenta, gold, green and orange, respectively). We hypothesize that these loops likely interact with the host’s surface and may be host-range determining regions (HRDR).

**Figure 2:**
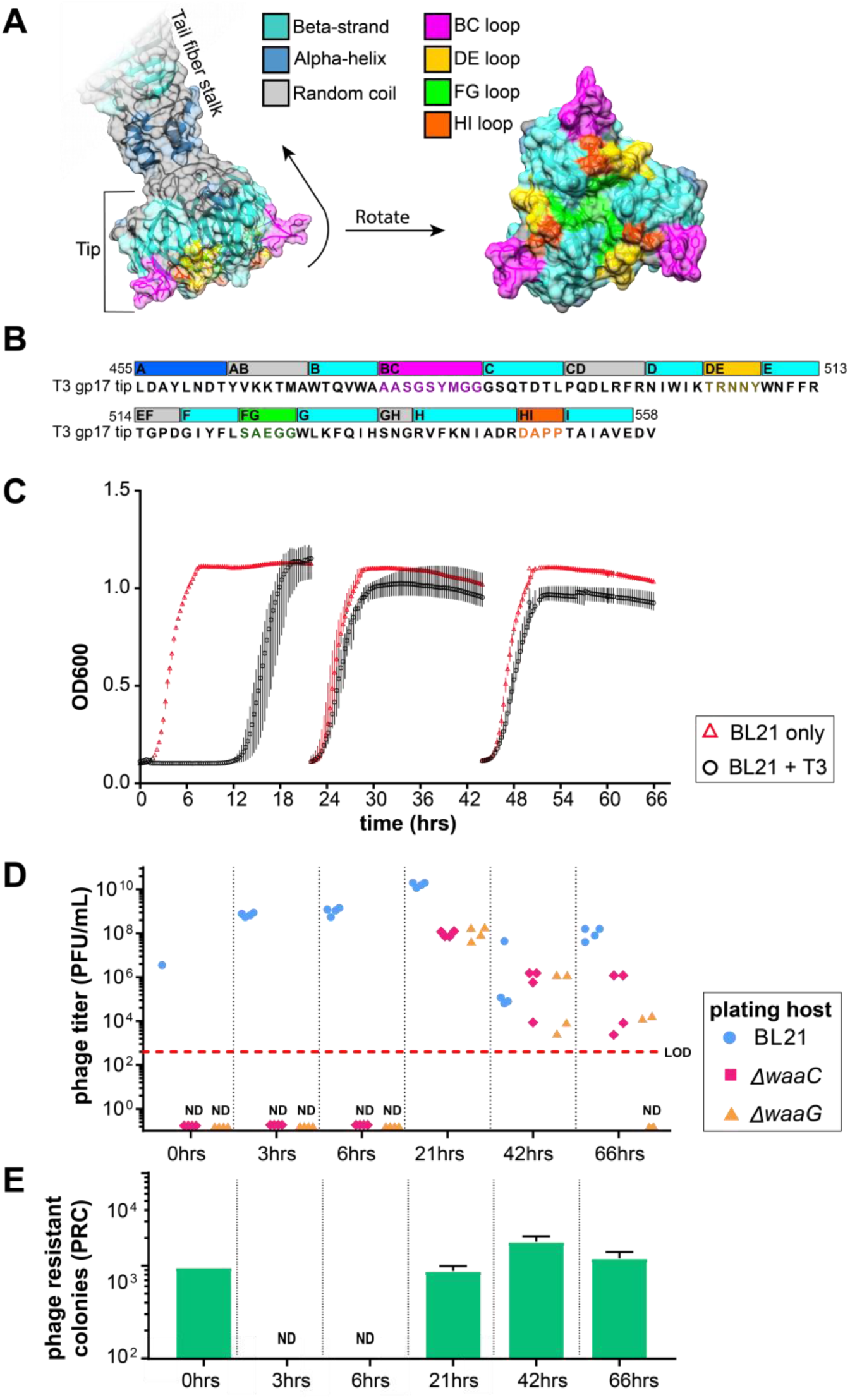
Computational structure of T3 gp17 and emergence of resistance to T3 infection. (**A**) Three-dimensional structure of the tail fiber tip domain of phage T3 as modeled by SWISS-MODEL. The molecular surface that include residues belonging to the BC, DE, FG and HI loops are highlighted in bright colors (magenta, green, gold, and orange, respectively) to illustrate their possible contribution to host binding. (**B**) Sequence of the T3 tail fiber tip (455-558 amino acids of the gp17 fragment) modeled in (**A**) with the same corresponding color scheme of structural features highlighted in the three-dimensional model. (**C**) Bacterial kinetic growth curves of BL21 on its own (red triangles) or BL21 infected with wild-type T3 (black circles). BL21 cultures were infected with wild-type T3 and reseeded every 24 hours into fresh LB medium. Data shown as the mean +/- standard deviation from three experiments. (**D**) Phage titers of the evolved T3 lysates on the parental host (BL21, blue circles) as well as on *ΔwaaG* (orange triangles) and *ΔwaaC* (red diamonds) were measured at the indicated timepoints. (**E**) At the indicated timepoints, ∼10^5^ plaque forming units (PFUs) of the evolved T3 lysates were mixed with ∼10^9^ wild-type BL21 colony forming units (CFUs) in a lawn. After 24 hrs of incubation at 37°C, the number of phage-resistant colonies (PRC) were counted (green bars; results presented as mean; error bars are standard deviation from three experiments; LOD: limit of detection; ND: not detected). T3 mutants that infect *ΔwaaG* and *ΔwaaC* appear during co-evolution with wild-type BL21, but these mutants in the evolved lysates are not capable of preventing resistance in culture or colonies from appearing in the plate resistance assay.

To corroborate structural identification of T3 HRDRs, natural phage mutants were isolated and sequenced to identify mutations that alter host-range. Natural phage mutants were selected for by infecting BL21 with T3 for up to 3 days with daily reseeding into fresh medium (Figure 2C). Phage mutants arise because T3 eliminates wild-type bacteria and selects for resistant bacterial mutants, which in turn, selects for T3 mutants. Based on sequencing and complementation assays, most bacterial mutants contain mutations in the LPS biosynthesis pathways that alter the primary receptor of T3. Because LPS mutations are the main evolutionary pathway for bacterial resistance against T3 infection, two LPS mutants of BL21 were generated by independently replacing two LPS synthesis genes with an apramycin resistance cassette: the BL21 *ΔwaaG* mutants lack the outer core of the LPS, including the glucose moiety that wild-type T3 uses as a receptor, and the BL21*ΔwaaC* mutants are nearly devoid of LPS (Figure 3B) (Heinrichs et al., 1998). Both strains contain truncated LPS and were used to detect phage mutants with expanded host-range by plaquing the “evolved” phage lysates (grown ∼24, 48, and 72 hours) on each bacterial lawn. Using these two strains, we could detect phage mutants after 24 hours of co-culture (Figure 2D, red and orange data points), while bacterial resistance developed ∼12 hours after infection (Figure 2C and E). Thirty-four plaques were isolated at random from BL21 *ΔwaaG* and *ΔwaaC* mutants and were submitted for Sanger sequencing of gene *17*. Out of the 34 phages isolated, only 10 contained unique combinations of mutations in gene *17*, of which, were comprised of 1 mutation in the BC loop, 1 mutation in the FG loop, and 4 mutations in the HI loop (Table S1). Surprisingly, despite a large proportion of phage mutants that can infect BL21 LPS mutants, the lysates failed to suppress bacterial resistance in wild-type BL21 as determined by culture turbidity (Figure 2C) and the number of phage resistant colonies (PRC) measured using a plate assay (Figure 2E), where phage lysates were mixed with BL21 top agar and poured onto LB plates and grown for 24 hours to select for bacterial mutants resistant to infection. Together, this suggests that phage evolution is too constrained to suppress resistance.

**Figure 3:**
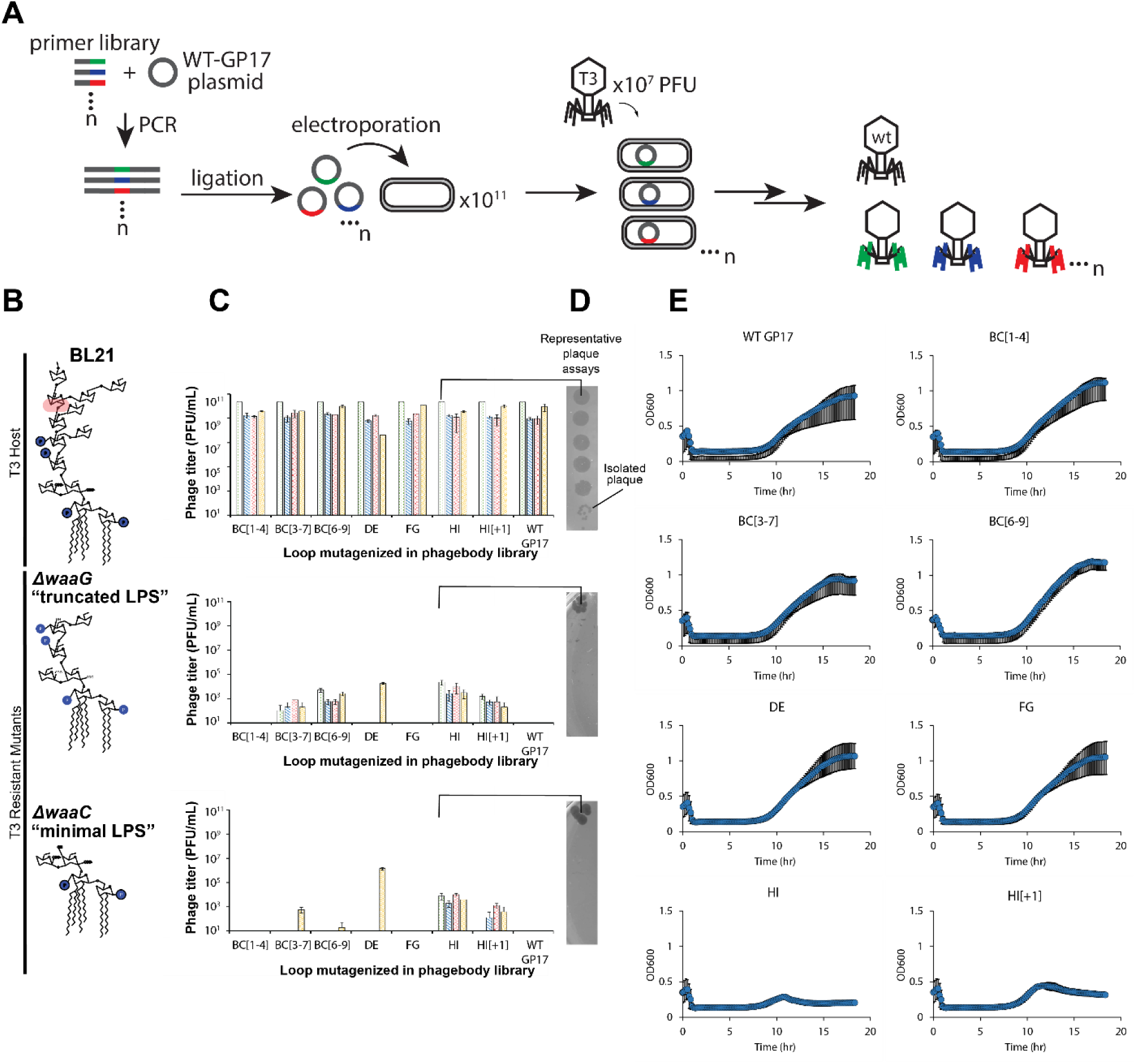
Phagebodies display broadened host-range towards LPS mutants that are resistant to wild-type T3 and can suppress bacterial resistance in culture. (**A**) Schematic showing the strategy to synthesize phagebody libraries. (**B**) LPS structures for wild-type BL21 and the T3-resistant BL21 mutants constructed for phagebody isolation. Highlighted in red is the sugar residue that is the T3 receptor. (**C**) Phage titers for 4 independent phagebody libraries (indicated by color) that had the indicated loop randomized. Titer was measured on wild-type BL21 (top row), *ΔwaaG* (middle row), and *ΔwaaC* (bottom row) in triplicate for each library and the data is plotted as mean +/- standard deviation. (**D**) Representative images of plaque assays from one of the HI loop phagebody libraries highlighting individual plaques. (**E**) Kinetic plots showing growth curves of wild-type BL21 bacterial cultures that were infected with phagebody libraries. As a control, wild-type T3 was grown on *E. coli* NEB5α carrying a wild-type T3 gene *17* plasmid (WT gene *17*) to account for non-specific mutations during library synthesis. Bacterial growth was monitored by measuring optical density at 600 nm. Each plot consists of 10 replicates from three independent experiments and shown as the mean +/- standard deviation. Cultures were infected at a MOI of 0.01.

### Strategy for introducing genetic variability in HRDRs

Because each of these loops are relatively small (4-9 a.a. long), we generated large diversity by replacing each codon within the targeted loops with NNK codons (Figure 3A), such that the loop sequences are completely randomized at the DNA level. To construct this library, we cloned the tail fiber gene *17* into a plasmid, which was PCR amplified using degenerate oligonucleotides designed to replace a single loop with a variable number of NNK codons. The resulting plasmid libraries containing mutated versions of gene *17* were transformed into *E. coli* NEB5α and recombined into the T3 genome by infecting transformants in early log-phase with T3. The infection did not proceed for longer than 3 hours to minimize enrichment of spontaneous phage mutants that arise from random mutations acquired by DNA polymerase errors (Figures 2C and E). Recombination with the plasmid was verified by measuring the ratio of the mutagenized gene *17* and the antibiotic-resistance gene (*kanR*) by qPCR. Silent mutations were introduced flanking gene *17* to selectively amplify the mutagenized gene *17* and not of WT using selective primers and a polymerase lacking 3’ to 5’ exonuclease activity (KAPA2G Robust) (Figure S1A). The ratio between gene *17* and *kanR* distinguishes between double recombinants versus single recombinants and nonspecific plasmid uptake (as a phagemid). Negligible amounts of the *kanR* gene was detected present inside the phage capsids (0.025%) (Figure S1B and C). The resulting recombined phages are referred to as phagebodies for the analogy to antibody specificity engineering.

Depending on when the plasmid-borne mutant of gene *17* is recombined in the phage genome, it is possible that both the wild-type gene *17* and the mutant version are concomitantly expressed, resulting in hybrid tail fibers (tail fibers are trimers of gp17) and chimeric particles that genetically encode a different gp17 than what is phenotypically assembled onto the virion particle. However, upon subsequent rounds of infection, phage progeny from these chimeras will fully express the encoded gene and no longer be chimeric. Therefore, we aimed for two rounds of infection to ensure expression of the mutated tail fiber, while minimizing library bias for enriching phages that infect BL21 (and not phage resistant mutants). This was achieved by infecting transformants at an MOI of 0.001, which equates to an average of 2 rounds of infection assuming a T3 burst size of 100. In addition, the majority of phages in the library are WT T3 because recombination efficiency is low (∼0.1-1%). Therefore, we included a control phagebody library (WT GP17) to account for any nonspecific mutations outside the targeted region that may impart host-range changes by infecting bacteria carrying an unmutated copy of gene *17*, resulting in only WT T3 phages.

The size of the theoretical sequence space for a given library mutating a single loop depends upon the number of codons mutagenized. For the smaller HI loop (4 codons), there are ∼10^6^ unique DNA sequences, while for the longest loop, BC (9 codons), there are ∼10^13^ unique sequences (NNK: 4×4×2^# of codons mutagenized^) (Table S2). Since it is not feasible to exhaustively sample the entire sequence space of the BC loop, we designed three separate libraries partially randomizing the BC loop, in which only the first four codons (BC[1-4]), the central five codons (BC[3-7]), or the last four codons (BC[6-9]) were randomized. The bracketed numbering indicates codon positioning within the loop (Figure S2A and B). In addition, libraries were generated that contained elongated HI loops compared to wild-type T3 with either one extra codon (HI[+1]) or three extra codons (HI[+3]). Figure 3A illustrates this pipeline.

### Characterizing phagebody libraries and screening on bacterial LPS mutants

To quantify library diversity and identify potential library construction biases, we performed NGS (HiSeq) at each step of library synthesis. Rarefaction curves for each sequenced library (Figure S2C) were plotted and showed that libraries targeting 4 codons saturated the theoretical DNA sequence space. Although libraries targeting 5 codons were not fully saturating at the DNA sequence level, they saturated protein sequence space when accounting for codon redundancy in the genetic code (Figure S2B and C). Libraries designed against loops longer than 5 codons were not saturating at either the DNA or protein level. Comparing the diversity differences between each stage in library construction (pre-transformed plasmid, transformed plasmid, and phage) suggests that the limiting step for library diversity is the transformation yield, while minimal loss in diversity can be attributed to recombination efficiency (Figure S2C). Thus, transformation yield was used as a measure to gauge library diversity.

To validate our hypothesis that loop randomization creates functional diversity, we screened phagebody libraries on *ΔwaaG* and *ΔwaaC* LPS mutants (Figure 3B), both of which are T3-resistant strains. Each phagebody library was serially diluted and arrayed on LPS mutant strains to quantify the number of PFUs and score the success of each library (Figure 3C and D). Library synthesis and screening were carried out multiple times to determine the reproducibility of library construction. A summary of all libraries constructed is available in Table S2. Screening of phagebody libraries on *ΔwaaG* and *ΔwaaC* bacterial mutants revealed that mutagenesis of particular loops are more amenable to broadening phage host-range than other loops, thus highlighting key phage-host interactions. Every library mutagenizing the HI loop produced phagebodies active against both *ΔwaaG* and *ΔwaaC* LPS mutants (Figure 3C), despite the fact that some of these libraries were far from saturating and loop elongation may have unpredictable consequences on tail fiber structure (Table S2, HI[+3] line). In contrast, phagebodies for which DE and FG loops were mutagenized, rarely demonstrated infectivity against *ΔwaaG* or *ΔwaaC* LPS mutants (Figure 3C; Table S2). Finally, approximately half of the libraries aimed at all or parts of the BC loop produced phagebodies capable of plaquing on *ΔwaaG* or *ΔwaaC* (Figure 3C; Table S2). Therefore, we conclude that mutagenesis of HI and BC loops can yield productive phage mutants, while the DE and FG loops are constrained in sequence by the tail-fiber structure or do not play a role in receptor binding.

### Phagebodies can delay or prevent bacterial resistance by loop mutations

Since phagebodies can plaque on and infect bacterial LPS mutants known to evolve from prolonged T3 infection (Perry et al., 2015), we tested whether these phage libraries can curtail resistance compared to wild-type T3. As an initial screen, we measured bacterial growth kinetics upon phage infection (MOI ∼0.01). Only the HI targeted libraries prevented bacterial resistance 24 hours post infection (Figure 3E). Surprisingly, the BC loop libraries did not curtail resistance despite the presence of LPS mutant-targeting phagebodies. Phagebody libraries are composed of a large diversity of unique phages (∼10^5^-10^7^ unique engineered phages), such that each individual phagebody is at a low concentration. Additionally, an unknown fraction of the phagebodies may not be functional or at least do not recognize the host upon which they are selected. To alleviate this constraint, we tested whether serial panning and amplification would uncover rare or poorly growing phagebodies. To implement this, three different mutant host strains were used, *ΔwaaG, ΔwaaC*, and a natural T3-resistant BL21 mutant that was isolated from a T3-resistant BL21 co-culture, referred to as PRM01 (<u>P</u>hage <u>R</u>esistant <u>M</u>utant). The panning experiment consisted of infecting a fresh culture of the desired bacterial strain with a phagebody library at a high multiplicity of infection (MOI=0.4), then recovering the progeny phages and repeating the cycle 3 times at lower MOI (Figure 4A).

**Figure 4:**
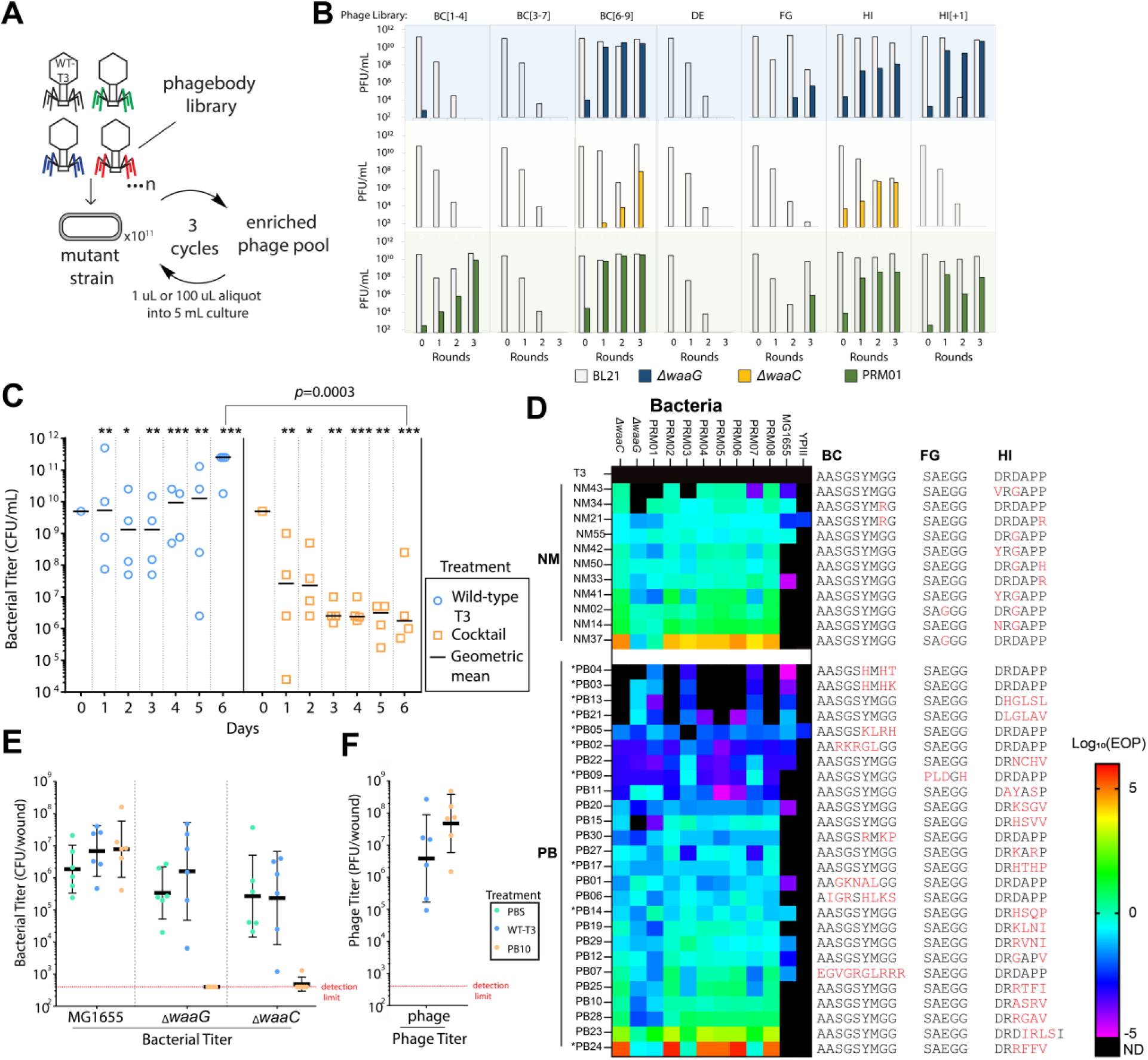
*In vitro* and *in vivo* activity of select phagebodies. (**A**) Schematic showing the phage panning procedure to amplify functional phagebodies from libraries. (**B**) Plots summarizing phage titers showing the amplification of functional phages and dilution of wild-type T3 per round of passaging on mutant strain. Rows are organized by the strain the library was passaged on: blue: *ΔwaaG*; yellow: *ΔwaaC*; and green: T3-resistant mutant PRM01 isolated from a wild-type BL21 culture infected with wild-type T3. Columns are organized by the phagebody library that was being passaged. (**C**) Replicates of four 50 ml cultures were inoculated with wild-type T3 (blue circles) or a cocktail of 10 phagebodies (orange squares) obtained from the enrichment experiment. Each culture was serially passaged every day with a 2-fold dilution into 2x-concentrated LB media for 6 consecutive days and the bacterial titer was measured each day. The day 0 titer corresponds to the starter culture before phage addition. All data points are represented with the geometric mean as a black horizontal bar. Each day shows statistically significant differences between the bacterial titers for cultures treated with the cocktail versus wild-type T3 (* *p*< 0.1, ** *p*<0.05, and *** p<0.001; one-tailed t-test on log-transformed data). The limit of detection is ∼300 CFU/ml, which is below the lowest data point on the graph. (**D**) Heat map summarizing the efficiency of plating (EOP; ratio of PFU on the bacterial mutant versus on wild-type BL21) for randomly isolated NMs and PBs on the two constructed isolation hosts, *ΔwaaC* and *ΔwaaG*, a panel of experimentally isolated wild-type-T3-resistant bacterial mutants PRM01-08, *E. coli* MG1655, and *Yersinia pseudotuberculosis* YPIII (ND: not detected; * indicates phagebodies used in the defined cocktail in **C**, which were isolated by enrichment on T3 resistant mutants). (**E**) Mice were infected with a consortia of bacteria that include equal amounts of MG1655, *ΔwaaC* and *ΔwaaG* (∼3×10^9^ CFU each) and were treated one hour post infection with PBS, T3 (∼10^9^ PFU) or PB10 (∼10^9^ PFU) (*n*=6 mice per treatment). 24 hours later, wounds were surgically removed and homogenized for colony counts and (**F**) plaque counts on BL21. The horizontal bar represents geometric mean and error bars show +/- standard deviation.

Despite amplification, the BC[3-7] and DE libraries still failed to produce phagebodies capable of infecting either of the three test strains. Phagebodies from the FG library were detected, but required an initial MOI = 40 (Figure 4B). Interestingly, FG libraries were also the most difficult to build, as they consistently yielded lower transformation efficiencies and had lower diversity than the other libraries (Table S2). A detailed analysis of NGS sequence distributions showed that FG libraries had a higher amount of WT T3 phage and even lower sequence diversity compared to other libraries (Figure S2D). The remaining phagebody libraries easily produced large titers of phagebodies that could infect T3-resistant strains with limited to no enrichment required (Figure 4B).

We next tested whether a defined cocktail composed of a limited number of phagebodies enriched from the panning experiment could curtail resistance similar to the HI targeting libraries. As an initial cocktail formulation, 10 isolated phagebodies from the panning experiments were mixed together in equal amounts (3×10^6^ PFUs each). Four of the phagebodies used in the cocktail had mutations in the BC loop, one had a mutation in the FG loop, and the remaining five had mutations in the HI loop (Table S3). Replicate 50 mL cultures of BL21 were infected with phage T3 or the cocktail at an MOI of 10^-3^ (total phage concentration: 3×10^7^ PFUs). Every 24 hours, each co-culture was diluted two-fold into fresh 2x LB medium. The low dilution rate was used to minimize dilution of any particular member of the phage cocktail that may be important for targeting bacterial mutants later in the infection, while also providing opportunity for bacteria to replicate and accumulate mutations that can confer phage resistance.

The cocktail-treated samples showed a decreasing trend in bacterial titer over time (Figure 4C, orange squares), while the T3-infected cultures rebounded to saturation after ∼12 hrs and maintained a constant bacterial titer of ∼10^10^ CFU/mL over the six days (Figure 4C, blue circles). The cocktail-treated cultures significantly outperformed the T3-treated cultures on each day using a single tailed t-test analysis on log-transformed data (Figure 4C, blue circles versus orange squares). By day 6, the largest difference in bacterial load was observed between cocktail-treated and T3-treated cultures, in which the phagebody-cocktail-treated samples achieved approximately 5 orders of magnitude reduction in bacterial titer compared to T3 (*p* = 0.0003) (Figure 4C, blue circles versus orange squares). As a result, the defined phagebody cocktail demonstrated similar capabilities to HI libraries, *albeit* with much lower genetic diversity in the phage population, and was capable of preventing regrowth of bacterial cultures for about a week. In contrast, T3 incurred visible resistance after ∼12 hrs and never managed to mount an effective response to control the bacterial population.

### Comparison between naturally evolved phages and phages with engineered HRDRs

To better understand how phagebodies are able to suppress bacterial resistance and whether there is a dependence on particular mutations or loop properties, we isolated additional phagebodies (PB) and naturally occurring mutants of T3 (NM) to compare their genomes, host-range, and killing efficiency. Host-range was determined by plaquing PBs and NMs on a panel of BL21 mutants, *E. coli* MG1655, and *Yersinia pseudotuberculosis* YPIII. The latter two strains were included to determine to what extent phagebody engineering expands host-range since it is known that T3 can evolve infectivity against these two strains (Garcia-Doval and van Raaij, 2012). The BL21 panel included the original three isolation strains (*ΔwaaG*, *ΔwaaC*, and PRM01) complemented with seven additional randomly isolated mutants of BL21 that acquired T3 resistance (PRM02-PRM08). Figure 4D summarizes the EOP host-range for PBs and NMs using their titer on BL21 as a reference, where Table S4 summarizes corresponding PFU counts. The results are presented as a heatmap (Log_10_EOP), where color indicates how well a phage performed on a given strain relative to BL21 and phage grouping is based on phage origin and performance (average EOP). As seen in Figure 4D, most NMs displayed a broad host-range for the BL21 bacterial panel, where 8/11 NMs were capable of infecting all 10 strains with similar efficiency as infecting WT BL21 (−2<Log_10_EOP<2). NM37 (FG: E525G) exhibited host switching by plaquing poorly on BL21, thus leading to a Log_10_EOP≥2 for most T3-resistant strains except for *Δ_waaG_* and PRM001. Interestingly, NM02 had the same mutation as NM37 (FG: E525G), but also contained an additional mutation (HI: D547G) that restores the ability to plaque on WT BL21. Lastly, 5/11 NM were able to infect MG16555, where only NM21 was capable of infecting YPIII.

On the other hand, PBs exhibited a more diverse range of host-range phenotypes. For example, for the BL21 bacterial panel, 4/26 PBs (PB4, PB3, PB13, and PB21) only infected a few strains (<6 strains) and at low efficiency (Log_10_EOP<-2); 6/26 PBs (PB5, PB2, PB22, PB9, PB11, and PB20) had broad host-range (≥9 strains) but also at poor effiency (Log_10_EOP<-2); 14/26 PBs infected all BL21 strains of the panel at efficiencies similar to infecting WT BL21 (−2≤Log_10_EOP≤2); and 2/26 PBs (PB23 and PB24) demonstrated host switching (Log_10_EOP>2), similar to NM37 but containing mutations in the HI loop rather than the FG loop. Similar to NMs, PBs were able to infect MG1655 more readily than YPIII, where 13/26 PBs could infect MG1655 and only PB05 could infect YPIII. This diversity in host-range phenotypes (narrow to broad) enables probing into how the physiochemical properties of each loop region correlates with host-range. Charge, hydrophilicity, and hydrophobicity were calculated for each peptide sequence of all phage mutants and correlation plots were generated by comparing the loop’s physiochemical properties with a phage’s host-range, as determined by the number of strains a phage is able to infect at Log_10_EOP>0.1. This analysis was performed for individual BC and HI loops, as well as, for all loops combined (Figure S3). Although, there were not enough phage mutants to generate FG loop correlation plots, these mutants are included in the combined analysis of all loops. Hydrophilicity and positive charge in the HI loop have the strongest correlation with broadened host-range (R^2^= 0.66 and 0.57, respectively) (Figure S3).

Next, the ability for NMs and PBs to suppress resistance was quantified using a plating assay, where ∼10^5^ PFU of phage were mixed with ∼10^9^ CFU BL21 in soft agar and poured over an LB plate and grown for 24 hours. After 24 hours, the number of phage resistant colonies (PRC) were counted and normalized to the number of PRC from T3 WT infection (Figure S4A). Typically, phages that infected all panel strains efficiently, which include PB10, PB12, or NM33, suppressed BL21 mutants that survived T3 infection (>99.9% killing of T3-resistant mutants). In fact, PB12 and NM33 left no detectable colonies after 24 hours. On the other end of the spectrum, phages that plaqued poorly on BL21, such as PB23, PB24, and NM37, failed to eliminate BL21, and therefore selected for high PRCs. In between these two extremes, the link between host-range and killing efficiency was only qualitative. For example, PB22 had broad host-range but poor infectivity towards any given PRM, but managed to eliminate 94.8% of the T3-surviving colonies. While PB9, despite having a very similar host-range pattern, allowed survival of 70% more colonies than T3. Overall, phage host-range only qualitatively correlates with how well a phage suppresses resistance (Figure S4B; R^2^∼0.5). However, interestingly, hydrophilicity of the HI loop correlates stongly (R^2^ ∼ 0.7) with resistance suppression (Figure S4B).

### PB10 retains specificity and is active in a mouse skin infection model

To test whether engineered phagebodies are functional *in vivo* and maintain selectivity, we tested the ability of phagebody PB10 to selectively kill sensitive bacterial strains in a consortia of bacteria using a mouse skin infection model. We chose PB10 because it showed broad infectivity against PRMs and had a hydrophilic HI loop, thus was anticipated it to prevent resistance development. Anesthetized mice were given a scratch (1 cm^2^ wound) and infected with a mixture of MG1655, *ΔwaaC* and *ΔwaaG* in equal amounts (∼3×10^9^ CFU each). One-hour post infection, wounds were treated with PBS, T3 (∼10^9^ PFU) or PB10 (∼10^9^ PFU). 24 hours later, wounds were surgically removed and homogenized for colony and plaque counts. The three bacterial strains could be differentiated based on their antibiotic resistance patterns (see Methods section). MG1655, which is resistant to both T3 and PB10, was used as a control to test whether phages became more promiscuous during wound treatment.

As expected, the MG1655 titer in the infected wounds did not vary significantly based on which treatment was applied (Figure 4E; ∼10^6^-10^7^ CFU/wound after 24 hours). Mice treated with PBS or T3 showed no significant decrease in *ΔwaaG* or *ΔwaaC* counts. On the other hand, 6/6 mice treated with PB10 had no countable colonies for *ΔwaaG* (Figure 4E; limit of detection ∼300 CFU/wound) and 5/6 mice had no countable colonies for *ΔwaaC* (Figure 4E; limit of detection ∼300 CFU/wound), where one mouse had a very low *ΔwaaC* count. Since PB10 yielded approximately an order of magnitude greater PFU counts (6×10^7^ PFU/wound) compared to T3 treated mice, determined by plaquing on BL21 (Figure 4E, right column), we conclude that PB10 was able to replicate and clear *ΔwaaG* and *ΔwaaC* from the wounds. Importantly, these results illustrate that PBs remain selective and active in animal tissues.

## DISCUSSION

In this paper, we present a powerful yet simple method to produce and screen libraries of engineered phages with expanded host-range and show that these engineered phages are able to suppress bacterial resistance to phage infection. We demonstrate that these engineered phages function similarly to NMs and can greatly scale the number of effective phages at preventing resistance, yet are composed of defined mutations. We also show that a limited cocktail of engineered phages can also suppress resistance, alleviating potential regulatory concerns of a phage library. Our directed approach relies on extensive mutagenesis targeting small regions of the phage tail fiber gene, which were selected based on available structural data for the likelihood to partake in host recognition. Our strategy is inspired by antibody specificity engineering, whereby (*i*) structurally important sequences are left intact, (*ii*) regions involved in target recognition are mutagenized, and (*iii*) the resulting libraries are screened against selected antigens (Ducancel and Muller, 2012).

Traditionally, new host-ranges can be achieved through natural evolution or through rational genetic engineering by swapping genes. However, the latter strategy requires prior knowledge of related genetic elements with the desired host-range and the capacity to engineer those elements into a functional phage, which often results in laborious *ad hoc* trial and error periods (Ando et al., 2015; Chen et al., 2017; Gebhart et al., 2017; Hawkins et al., 2008; Heilpern and Waldor, 2003; Lin et al., 2012; Nguyen et al., 2012; Scholl et al., 2009; Yoichi et al., 2005). Harnessing natural evolution, on the other hand, requires elaborate co-culture and selection strategies that balance the need to have the phage population grow and mutate while also providing an alternative host for phage mutants to reproduce on (Qimron et al., 2006; Ross et al., 2016; Yu et al., 2015). A recent paper by Yosef and colleagues combined the approaches of rational host-range engineering with evolution. In particular, they engineered phages that lack a tail fiber gene and provided the missing function from a library of plasmids encoding various tail fiber homologs from phages with known host-range. The tail-encoding plasmids were then evolved through chemical mutagenesis and selected based on the capacity for phages to transduce the corresponding plasmid to a host (Yosef et al., 2017).

In contrast, the approach described herein focuses on producing viable phages with subtle host-range alterations to target resistant mutants. We directed the mutagenesis to regions of the tail fiber that are expected to be most productive, in host-range determining regions (HRDR). Because these regions are short, we can saturate the potential sequence space and produce large libraries of phages with diverse functionalities that range from more specific to broadened host-range. Moreover, we can discover phage mutants that would require too many point mutations to be readily produced by artificial evolution. Importantly, because of the diverse host-range phenotypes generated, we are able to determine key phage-host interactions critical for suppressing resistance. We expect this method will greatly aid in mapping key residues for receptor recognition and determining which amino-acid substitutions lead to host-range alterations, thus providing a rich tool for structural biology of phage receptor recognition. Additionally, if the structure of the phage protein is unknown, our method is rapid and simple enough that it can be used to scan for the most important regions involved in host recognition, as was exemplified by our work on the BC loop. If the tail fiber structure is unavailable, these regions can be identified through NGS analysis of evolved phage lysates.

Furthermore, using cocktails of natural phages can be challenging from a translational perspective because of the need to manufacture diverse phages, whereas phagebody cocktails can be based off of a common scaffold. In addition, phagebodies can be further enhanced by selection for other phenotypes (e.g., replication rates or altered immunogenicity) and natural evolution can be used to enrich commensurate mutations outside of the immediately targeted region. Yet, our approach is simple enough that iterative cycles can be performed to generate phagebodies mutating several loops concurrently.

However, resistance to phage infection does not exclusively arise from receptor mutations. For example, abortive infection systems, CRISPR systems, and encapsulation by exopolysaccharide that prevents receptor access are common resistant mechanisms. Further work is required to devise strategies to overcome these pathways, which we think is an exciting direction for phage engineering. For example, abortive infection systems are often triggered by phage activity or a peptide sequence. When such triggers are known, they can be engineered out of the phage to render them insensitive to these resistance mechanisms (Labrie et al., 2010). Alternatively, various phages have evolved CRISPR inactivating proteins (Pawluk et al., 2016), which can be cloned into our scaffold bacteriophage. Phages expressing capsule-degrading enzymes and using capsules as receptors are not uncommon (Hsu et al., 2013; Kim et al.; Leiman et al., 2007; Lin et al., 2014; Pickard et al., 2008). Therefore, it is possible to engineer capsule-degrading catalytic activity into phages that lack this function (Gladstone et al., 2012; Lu and Collins, 2007). Alternatively, instead of relying on phage replication and antimicrobial activity, one can use engineered phage capsids to deliver therapeutic payloads into bacteria, such as CRISPR-Cas nucleases (Bikard et al., 2014; Citorik et al., 2014).

We envision that the high-throughput mutagenesis and characterization of host-range determinants will be an important and useful tool to decipher host recognition by viruses. As more and more viral tail components are structurally resolved and sequenced (Montag et al., 1987; Pouillot et al., 2010; Tétart et al., 1996; Trojet et al., 2011), the breadth of viral models on which this strategy can be applied will also be expanded. Moreover, mapping phage-host interactions should enable the rational engineering of phage host-range. The speed and scale at which phagebodies can be engineered and screened should enable rapid adaptation and discovery of effective antimicrobial agents against bacterial pathogens. Finally, the ability to discover and optimize novel antimicrobial agents that can achieve long-term suppression of bacterial resistance has the potential to significantly improve treatment of AMR.

## EXPERIMENTAL PROCEDURES

### Strains and culture conditions

Bacteriophage T3 was obtained from Ian Molineux (University of Texas, Austin) and maintained on *E. coli* BL21. Cloning was performed in *E. coli* NEB5α. Bacteria were grown in Lysogeny Broth (LB medium; LabExpress) at 37°C with agitation at 250 rpm and stored at −80°C in 45% glycerol. As needed, the medium was supplemented with kanamycin (50 µg/ml final concentration), carbenicillin (50 µg/ml final concentration), apramycin (50 µg/ml final concentration), or glucose (0.2% w/v final concentration). LB plates contained agar (LabExpress) at a final concentration of 1.5%. Top agar is LB agar at 0.6%. Bacterial strains resistant to wild-type T3, PRM001-PRM008, were picked from wild-type-T3-infected lawns of wild-type *E. coli* BL21 incubated at 37°C until resistant colonies grew. They were picked, streaked to isolation twice, and tested for T3 resistance.

### DNA manipulation and cloning

PCR was performed using either KAPA Biosystems HiFi or Robust 2G DNA polymerases using recommended PCR conditions for each polymerase. DNA was purified using Zymo Research reagents. Restriction enzymes were purchased from New England Biolabs. All reagents and kits were used following the manufacturer’s recommendations.

pSLM49 was constructed by assembling the PCR-amplified replication origin and resistance marker of pFF753 (primers PST480 and PST481) (Farzadfard and Lu, 2014) with a PCR-amplified fragment from phage T3 containing gene *17* (PST575 and PST576) through BamHI/XmaI restriction cloning. pSLM193-197 and pSLM225-233 are derivatives of pSLM49 built by cloning the gene *17* tip sequence from select phagebodies in lieu of the wild-type tip sequence. The gene *17* tips were amplified through primers PST691/692 and the rest of the plasmid with PST693/694. The two PCR fragments were then assembled by Gibson® reaction. pSLD18 is a derivative of pSIM9 (Datta et al., 2006) where the chloramphenicol marker was replaced with the erythromycin marker of pCP1 (Le Bourgeois et al., 1992). pSLM111alpha was obtained by ligating the apramycin resistance marker of plasmid pSET152 (Bierman et al., 1992) amplified with primers PST816 and PST817 and the backbone of pKD3 (Datsenko and Wanner, 2000) amplified with primers PST818 and PST819 after restriction of both fragments with PspoMI.

### LPS mutant construction

*E. coli* BL21 was transformed with the recombineering plasmid pSLD18 and cells were made recombineering proficient as described in (Datta et al., 2006). The cells were electroporated with a PCR product designed to replace *waaC* or *waaG* with an apramycin resistance marker amplified from pSLM111alpha with primers PST853/PST854 and PST857/PST858, respectively. Proper deletion was verified by PCR.

### Tail fiber library creation

Diversity was introduced at the DNA level in pSLM49 using two different methods: (1) Direct transformation of PCR products with terminal redundancy and (2) a restriction-ligation based method.

#### 1. Direct transformation of PCR products

The entire pSLM49 plasmid was PCR amplified with a pair of diverging primers annealing on each site of the target loop. In one of the oligonucleotides, the target loop sequence was replaced by a series of NNK codons. The NNK stretch was preceded at the 5’ by the complementary sequence to the reverse primer so that the final PCR product had a 20-30 bp identical sequence at each end. The amplicons were then DpnI digested to eliminate template DNA and about 100-500 ng of the resulting DNA was transformed into chemically competent NEB5α following the manufacturer’s instructions. The termini of the PCR products were redundant such that the PCR product was circularized, reconstituting gene *17* but with a random sequence in place of the targeted loop. The bacteria were recovered for 1 hour at 37°C in SOCS medium (1 mL). After this step, the transformation yield was determined by plating serial dilutions of culture on LB-kanamycin agar plates. The 1 mL bacterial cultures were then diluted with 9 mL of LB-kanamycin and grown overnight at 37°C and 250 rpm of shaking. The next day, fresh cultures were started by diluting 1 mL of overnight culture into 9 mL of LB, while the remaining culture was pelleted and stored at −20°C for plasmid DNA extraction/sequencing. Phage lysates were made by infecting bacterial cultures at exponential growth phase (OD_600_= 0.7) with 10^7^ plaque forming units (PFU) of T3 (100 µL). The cultures were grown for another 2-3 hours until the cultures cleared. Phage lysates were chloroform treated with 500 µL of chloroform for 30 minutes to kill any remaining bacteria, spun down to remove debris and filtered through a 0.22 µm filter. Phage lysates were spun down at 7,000 G for 5 minutes and stored at 8°C for long-term storage.

#### 2. Restriction-ligation-based method

The entire pSLM49 plasmid was PCR amplified with a pair of diverging primers annealing on each site of the targeted loop using HiFi polymerase following manufacturer recommended protocol. Primers were designed to encode (1) a mutagenized region corresponding to the desired *gp17* loop and (2) BsaI cleavage sites for restriction digestion, which was digested and subsequently circularized by T4 ligase to yield libraries of scarless circular plasmids (Figure 3A). The mutagenized region was encoded by NNK codons to minimize premature incorporation of stop codons.

Each PCR reaction was pooled together and DpnI digested. Following digestion, the PCR products were purified using Zymo DNA clean and concentrator^TM^-5 spin columns. Next, ∼5 µg of linearized *gp17* gene products were diluted in New England Biolabs CutSmart® buffer (500 µL) and restriction digested using 125 units of BsaI at 37 °C for four hours. The enzyme was heat inactivated at 65°C for 20 minutes. Digested DNA was purified using Zymo DNA clean and concentrator^TM^-5 spin columns and eluted in Nanopure water (18.2 MΩ).

The digested DNA was circularized using T4 ligase, where 2 µg of DNA was diluted to 50 µL in T4 ligase buffer and 1 µL of T4 ligase (400,000 units) was added. The reaction was incubated overnight at room temperature. The next day, DNA was purified using the Zymo DNA clean and concentrator^TM^-5 spin columns and eluted with 10 µL of Nanopure water to yield a plasmid wild-type of ∼100 ng/µL. Next, bacterial libraries were made by transforming 100 ng of plasmid into New England Biolabs NEB5α electrocompetent cells via electroporation (1 mm cuvette, 1.7 kV, 200 Ω, and 20 µF). The transformed bacteria were then handled as described for method 1.

### qPCR analysis

KAPA SYBR® FAST qPCR Kit was used to quantify the number of mutated gene *17* and *kanR* genes. Calibration curves were made by varying the amount of plasmid encoding both gene *17* and *kanR* (x) and measuring Cq (y: cycle of detectable fluorescence) for both genes. Phagebody lysates were treated with TURBO DNase (Thermo Fisher Scientific) to degrade any soluble plasmid not encapsulated by the phage and then heat inactivated. pPCR was performed by measuring Cq for both genes and the copy number for each gene was calculated using the calibration curves. Importantly, SYBR®FAST polymerase does not have 3’-5’ exonuclease activity, which is important for selectively amplifying the mutant version of gene *17* and not wild type. The ratio between gene *17*:*kanR* is (x+y+c)/(y+c), where x is the number of phages that had double recombination, y is the number of phages that had single recombination, and c is the number of phages with full plasmid encapsulation. The measured ratio for all phage lysates was ∼4000:1. Solving for x (double recombination) results in x= 3999*(y+c), which is ∼4000×fold greater than the single recombination plus plasmid uptake.

### NGS sequencing and analysis

Illumina adapter sequences and barcodes were added to the plasmid DNA samples through overhang PCR using Kappa HiFi (25 cycles). For phage samples, phage lysates were treated with TURBO DNase I (Thermo Fisher: AM2238) according to manufacture protocol to eliminate any soluble plasmid. After which, phages were concentrated 50 fold using PEG precipitation and phage DNA was purified using Zymo’s Quick-DNA/RNA Viral Miniprep Purification Kit (D7020). To selectively amplify phagebody DNA versus WT T3 DNA, primers targeting silent mutations only present in the phagebody tail fiber gene were used to amplify 2 kb of the tail fiber region (25 cycles) using Kappa Robust 2G. WT gene *17* does not amplify because it contains 3’ mismatches (Figure S1A). Kappa Robust 2G was used over Kappa HiFi because Kappa HiFi has 3’→5’ exonuclease activity and will also amplify WT gene *17*. PCR amplicons were gel purified and the adapter sequences and barcodes were added through overhang PCR using Kappa HiFi (25 cycles). DNA was sequenced using Illumina HiSeq 2000 and 150 nt single end reads. The adapter and barcode sequences were trimmed using seqtk and rarefaction curves were generated by subsampling these data using seqtk and FASTAptamer (Alam et al., 2015) to quantify and identify unique sequences.

### Measuring efficiency of plating (EOP) of phage lysates

Phagebody libraries were serially diluted in triplicate and 3 µl of each dilution were spotted onto the surface of 10×10 cm LB agar plates covered with a top agar lawn of the desired test strain. The EOP was calculated as the ratio between the phage titer on the mutant strain and the reference strain (*E. coli* BL21). A pseudocount of 1 was added to the entire dataset prior to any calculation.

### Phage panning

For each bacterial mutant, overnight cultures were grown from a single colony. The next day, 50 µL of the overnight culture was diluted into 5 mL of LB and grown to exponential phase (OD_600_: 0.7), at which 100 µL of phage lysate was added. The bacterial cultures were grown for another 3 hours, except for *ΔwaaC* mutants, which the culture was grown for 4 hours. After phage propagation, phage lysates were chloroform treated (250 µL), spun down at 7,000 G for 5 minutes, and stored at 8 °C for subsequent panning. This procedure was repeated for additional rounds (Figure 4A and B), except infecting with 1 µL of phage lysate from the previous round of infection rather than 100 µL. This enabled amplification of functional phages, while diluting away phages incapable of infecting bacterial mutants.

### Resistance index determination

Triplicate samples of ∼10^5^ PFU of each phagebody isolate were mixed with ∼10^9^ CFU wild-type *E. coli* BL21 in 3 ml of top agar and immediately poured over LB plates. After the top agar hardened, plates were incubated for 24 hrs at 37°C. CFU were subsequently counted for each plate. Results were systematically normalized to the number of CFU counted on T3-infected lawns. A pseudocount of 1 was added to the entire dataset prior to any calculation.

### Liquid culture assay for resistance suppression by phagebody libraries

From an overnight culture of wild-type BL21, a fresh culture was grown to exponential phase (OD_600_= 0.7). After which, 250 µL aliquots of the culture were added to a 96 well plate along with 2.5 uL of phagebody lysates per well. This equates to an MOI of ∼0.01. Growth curves were obtained by taking OD_600_ measurements using a BioTek Synergy H1 microplate reader at 10 min. intervals, 37°C, and constant shaking.

### Liquid culture assay for resistance suppression by phagebody cocktail

From an overnight culture of BL21, 500 µL were diluted into 50 mL of LB in 250 Erlenmeyer flasks and grown to exponential phase (OD_600_= 0.7). Then, ∼10^7^ PFUs (MOI of 10^-3^) of phage lysate from wild type T3 (100 µL) or a phage library (10 µL) was added. The cultures were grown overnight. The next day, 1 mL aliquots were taken from each culture, washed 4 times in PBS, serially diluted and plated on LB-agar plates to quantify the number of CFU. Every 24 hours, the culture was diluted 2-fold with 25 mL of 2x-concentrated LB to ensure bacterial nutrients were still available.

### Scarification skin infection mouse model

All animal study protocols were approved by the MIT Animal Care and Use Committee. An overnight culture was transferred to eight 50 mL falcon tubes and spun down at 8,000G for 3 minutes at 4°C and washed with PBS. This step was repeated 4 times while concentrating bacteria ∼2 fold during each wash. For the final resuspension, bacteria were resuspended into 2 mL of PBS, which equates to ∼10^11^-10^12^ CFU/mL. To generate a skin infection, female CD-1 mice (6-weeks-old) were anesthetized with isoflurane and had 2 cm^2^ area of their backs shaved. A superficial linear skin abrasion (∼1 cm^2^) was made with a needle in order to damage the stratum corneum and upper-layer of the epidermis. Five minutes after wounding, an aliquot of 20 μL containing ∼10^10^ CFU of bacteria in PBS was inoculated over each defined area containing the scratch with a pipette tip. One hour after infection, 10^9^ PFU of phages were administered to the infected area. Animals were euthanized and the area of scarified skin was excised one-day post-infection, homogenized using a bead beater for 20 minutes (25 Hz), and serially diluted for CFU and PFU quantification. Experiments were performed with 6 mice per group. Statistical significance was assessed using a two-way ANOVA.

## AUTHOR CONTRIBUTIONS

KY and SL conceived of the concept and designed experiments; KY, SL, ACY and HA synthesized, characterized, and screened phage libraries; MT and CFN performed animal studies, MM helped with analysis, SL, KY, and TL wrote the paper.

## ACKNOWLEDGMENTS

We would like to thank Robert Citorik and Fahim Farzadfard for continued constructive discussions all along this project. Bacteriophage T3 was a kind gift of Ian Molineux, University of Texas, Austin. This work was supported by grants from the Defense Threat Reduction Agency (HDTRA1-14-1-0007), the National Institutes of Health (1DP2OD008435, 1P50GM098792, 1R01EB017755), and the U.S. Army Research Laboratory / Army Research Office via the Institute for Soldier Nanotechnologies, under contract number W911NF-13-D-0001. HA was supported by fellowships from the Japan Society for the Promotion of Science and the Naito Foundation. This work was supported in part by the Koch Institute Support (core) Grant P30-CA14051 from the National Cancer Institute.

---

**Figure S1:**
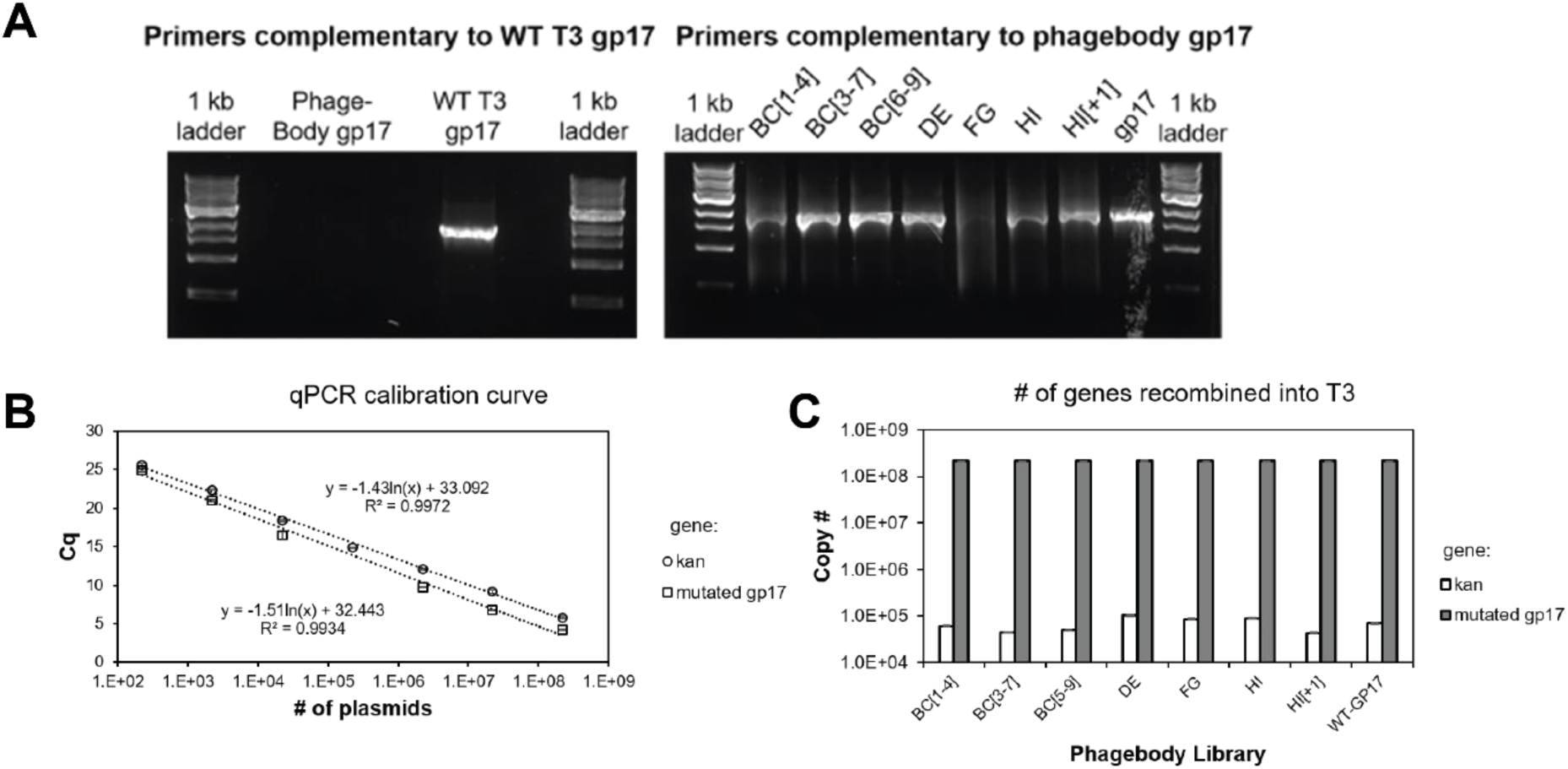
qPCR design and analysis of plasmid packaging into the phage. (**A**) The phagebody construct was designed to contain two 3 bp silent mutations at serine 71 in gene *16* and serine 25 in gene *17.5*. When using the appropriate primers, Kappa Robust 2G selectively amplifies either WT *17* (left) or phagebody *17* (right) depending on primers used. (**B**) Calibration curve used to quantify copy # of *kanR* and mutagenized *gp17* genes (Cq: quantification cycle). Error bars are +/- standard deviation of the mean calculated from 3 measurements. (**C**) Plot showing the amount of gene *17* and *kanR* in each phagebody library. The ratio between gene *17*:*kanR* is (x+y+c)/(y+c), where x is the number of phages that had double recombination, y is the number of phages that had single recombination, and c is the number of phages with full plasmid encapsulation. The measured ratio for all phage lysates was ∼4000:1. Solving for x (double recombination) results in x= 3999*(y+c), which is ∼4000×fold greater than single recombination plus plasmid uptake.

**Figure S2:**
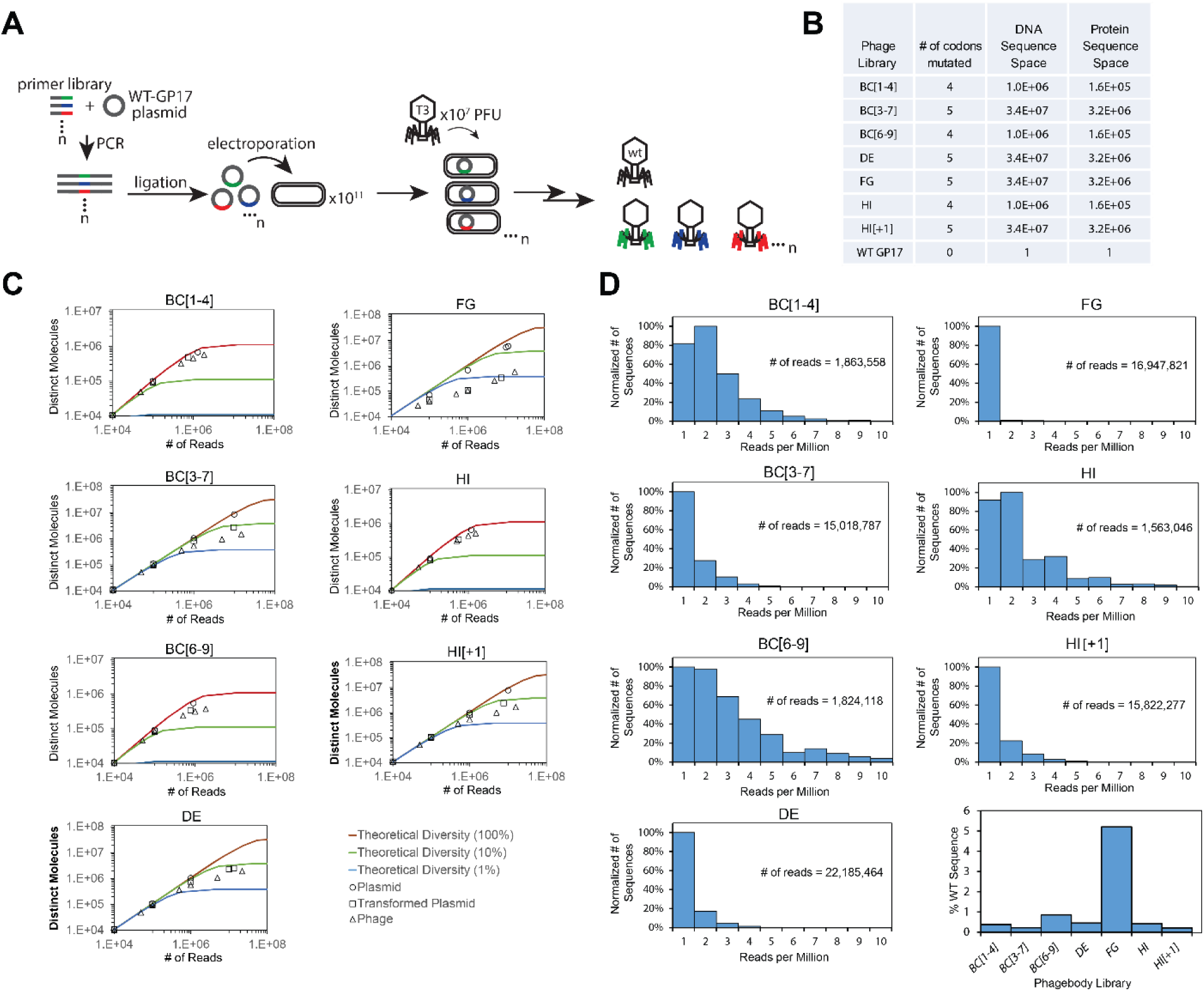
Illumina HiSeq characterization of phagebody libraries. (**A**) Schematic showing the strategy to synthesize phagebody libraries. (**B**) Table summarizing the theoretical diversity for libraries synthesized using Method 2 and corresponding library coverage calculated by (transformation yield/DNA library size) ×100. (**C**) Rarefaction curves characterizing library diversity at each stage of synthesis. Circles correspond to synthesized plasmid libraries, squares correspond to plasmid libraries recovered post transformation, and triangles correspond to phage libraries post recombination. Colored solid lines are guides showing 100% (red), 10% (green), and 1% (blue) coverage of theoretical sequence space (see calculation below). (**D**) Histogram analysis showing sequence distributions for each phagebody library. Each library had a single sequence at very high frequency. Upon further analysis, this sequence was wild type (WT) sequence. It is important to note that FG libraries had the largest amount of WT contamination and required multiple rounds of panning to isolate a phagebody. This indicates that phagebodies are much more dilute in FG libraries. Calculation for theoretical rarefaction curves: The theoretical diversity was calculated using the mathematical analysis derived from the classic “coupon collector” problem. Below is the equation for calculating the average number of reads required to capture diversity of a library. Average # of reads required to sequence entire library = (# of reads to identify 1st unique phage) + (# of reads to identify 2nd unique phage) +…. (# of reads to identify unique phage n) = n/n + n/(n-1) + …+ n/1 = n*[(1/1) +(1/2)+…(1/n)]

**Figure S3:**
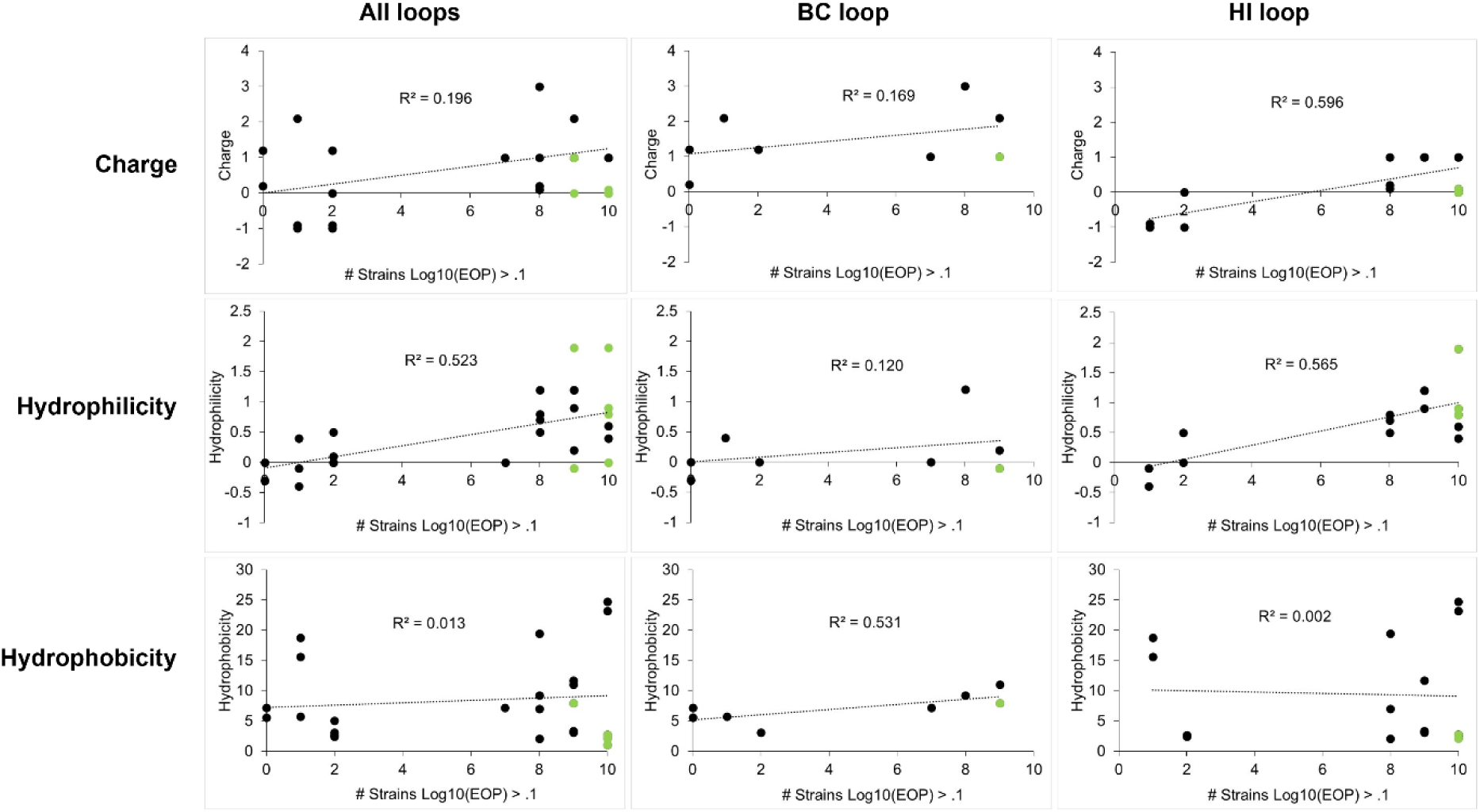
Host-range correlation plots comparing a loop’s physiochemical properties and phage host-range. Correlation plots comparing the loops’ physiochemical properties (top: charge; middle: hydrophilicity; and bottom: hydrophobicity) to phage host-range, as determined by the number bacterial strains a phage can infect at Log_10_EOP>0.1 from the BL21 T3 resistant bacterial panel in Figure 4D (PB, black circles; NMs, green circles). The lines show linear fit analyses with corresponding R^2^ values.

**Figure S4:**
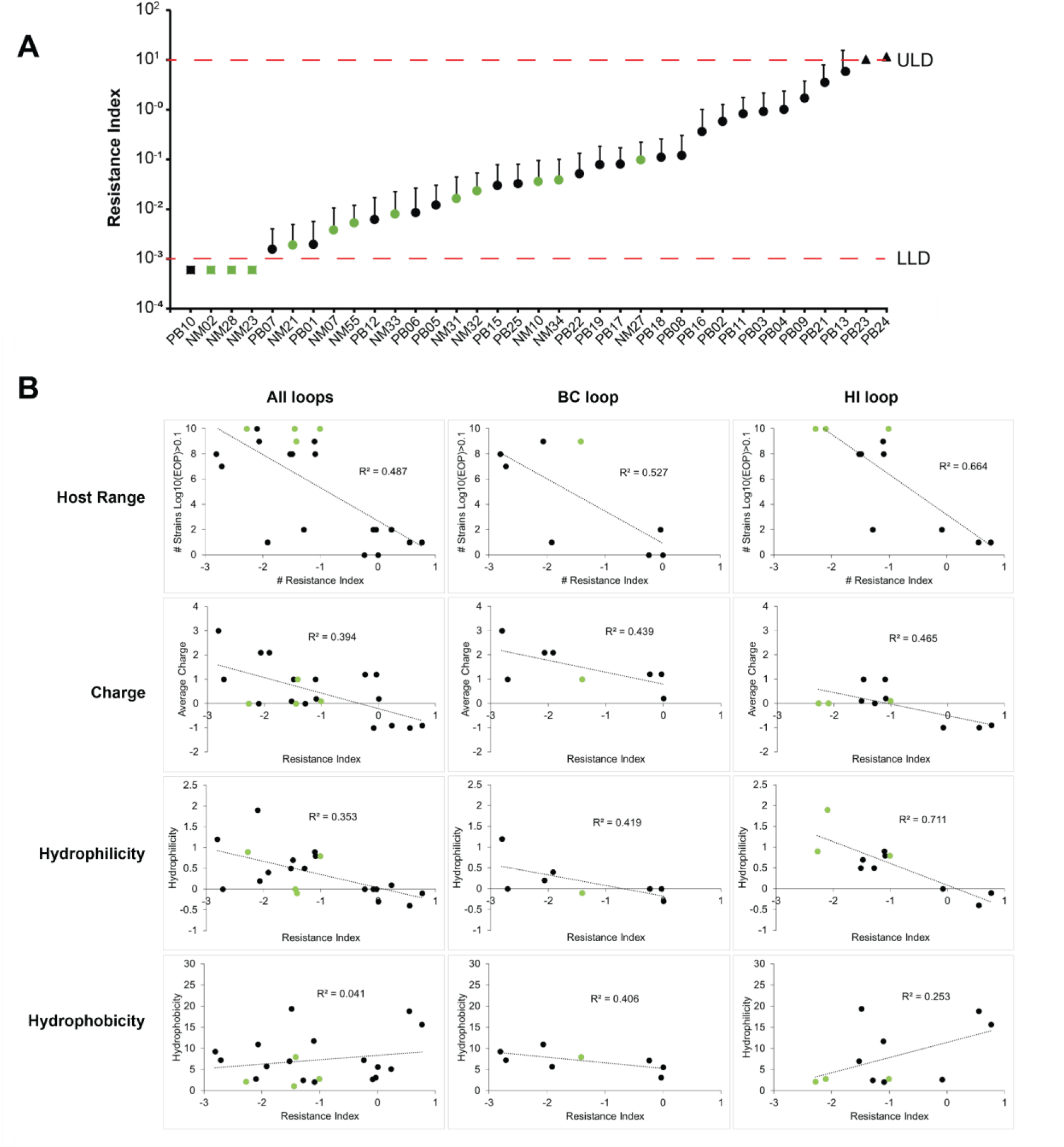
Resistance index comparing NMs and PBs ability to suppress resistance and correlation plots comparing phage host-range and loop physiochemical properties to resistance suppression. (**A**) A plot summarizing killing efficiency and resistance suppression for each phage mutant for both PBs (black) and NMs (green). ∼10^5^ PFU of phage were mixed with ∼10^9^ CFU BL21 in soft agar and poured over an LB plate and grown for 24 hours at 37°C. After 24 hours, the number of phage resistant colonies (PRC) were counted and normalized to the number of PRC from T3 WT infection. Resistance Index = Log_10_[(1+PRC_PB or NM_) / (1+PRC_WT-T3_)]. Error bars show upper standard deviation (Square: too few PRC to accurately count; triangle: too many PRC to accurately count; LLD: lower limit of detection; and ULD: upper limit of detection). (**B**) Correlation plots comparing resistance index with phage host-range (top) and loop charge (upper middle), hydrophilicity (bottom middle), and hydrophobicity (bottom) (PBs, black circles; NMs, green circles). The lines show linear fit analyses with corresponding R^2^ values.

**Table S1:**
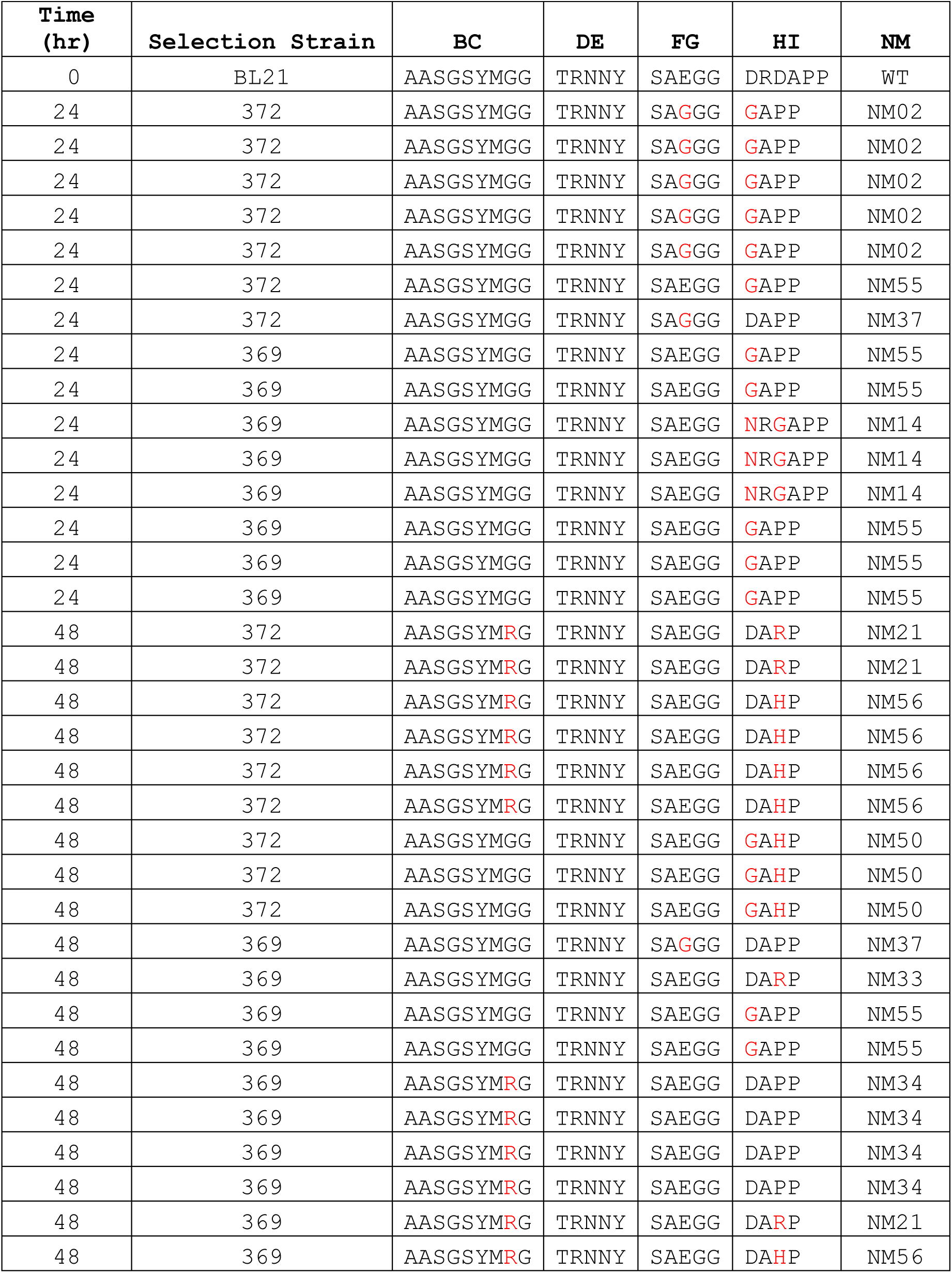
Summary of Sanger sequencing results for the gp17 tip region of natural T3 phage mutants isolated from 24 and 48 hr co-cultures of T3 and BL21. A BL21 culture (5 mL; OD = 0.7) was infected with T3 (10^7^ PFU) and was co-cultured for 24 and 48 hrs. After which, phage lysates were chloroform treated and filtered. Natural phage mutants were selected for and plaque purified on either isolation hosts: *ΔwaaC* or *ΔwaaG*.

**Table S2:** Cumulative summary of phagebody libraries constructed during this work. The theoretical diversity expresses the total number of possible DNA combinations based on the number of NNK codons randomized (4×4×2^# of codons mutagenized^). The cumulative coverage is the sum of the library transformation yields for all the libraries ever constructed for that loop. Calculated cumulative coverage is the percentage of theoretical diversity as determined by the total number of plasmid clones obtained for all repeats for each type of library. “Hits” are defined as obtaining at least one PFU on a lawn of the corresponding selective BL21 mutants, *ΔwaaC* or *ΔwaaG*

| Targeted loop | Theoretical diversity<br>(number of different<br>combinations from<br>NNK codon<br>randomization) | Number<br>of<br>libraries<br>built | Cumulative<br>coverage<br>(% of theoretical<br>diversity) | Number of<br>phagebody<br>libraries<br>producing<br>“hits” on<br>*ΔwaaC* | Number of<br>phagebody<br>libraries<br>producing<br>“hits” on<br>*ΔwaaG* |
| --- | --- | --- | --- | --- | --- |
| WT gp17 | NA | 21 | NA | 0 | 0 |
| BC | 3.5E+13 | 10 | ~0.0000005 | 4 | 7 |
| BC[1-4] | 1.0E+06 | 10 | ~60 | 4 | 6 |
| BC[3-7] | 3.4E+07 | 10 | ~90 | 2 | 5 |
| BC[6-9] | 1.0E+06 | 8 | >100 | 2 | 4 |
| DE | 3.4E+07 | 10 | ~50 | 2 | 4 |
| FG | 3.4E+07 | 10 | ~0.3 | 0 | 0 |
| HI | 1.0E+06 | 15 | >100 | 15 | 15 |
| HI[+1] | 3.4E+10 | 4 | ~0.00002 | 3 | 4 |
| HI[+3] | 3.4E+07 | 14 | ~20 | 10 | 10 |

**Table S3:**
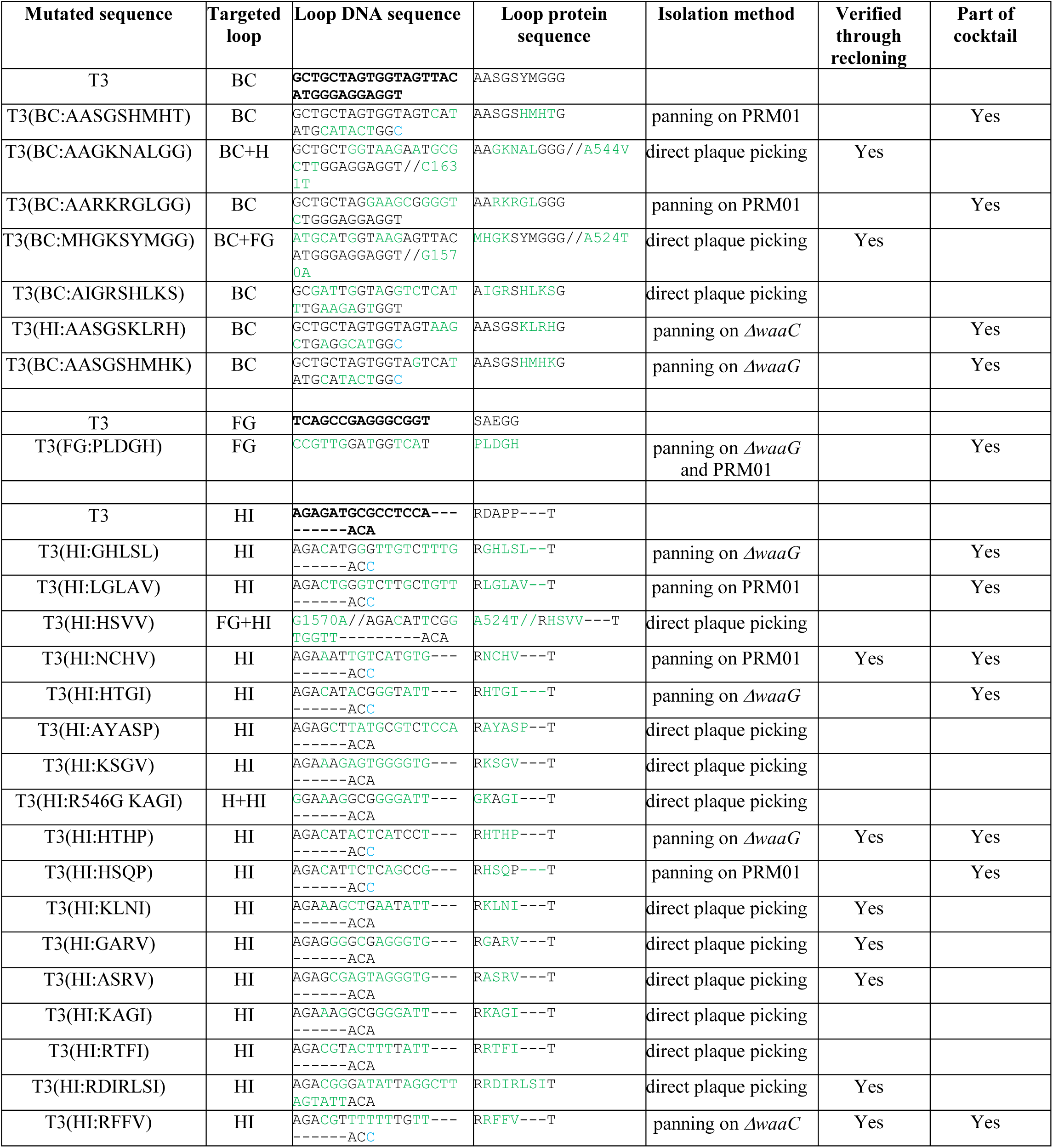
Summary of the phagebody strains characterized during this work. Summary of all the phagebodies isolated and characterized during this work showing mutated sequence, isolation method, whether the phagebody was verified through recloning the tip into T3, and if used in the minimal cocktail (Figure 4C).

(See attached spreadsheet- “EOP Summary”)

**Table S4. Summary of PFU counts used to construct EOP heatmap.**

